# High-fidelity base editor with no detectable genome-wide off-target effects

**DOI:** 10.1101/2020.02.07.939074

**Authors:** Erwei Zuo, Yidi Sun, Tanglong Yuan, Bingbing He, Changyang Zhou, Wenqin Ying, Jing Liu, Wu Wei, Rong Zeng, Yixue Li, Hui Yang

**Affiliations:** Center for Animal Genomics, Agricultural Genome Institute at Shenzhen, Chinese Academy of Agricultural Sciences, Shenzhen 518124, China; Institute of Neuroscience, State Key Laboratory of Neuroscience, Key Laboratory of Primate Neurobiology, CAS Center for Excellence in Brain Science and Intelligence Technology, Shanghai Research Center for Brain Science and Brain-Inspired Intelligence, Shanghai Institutes for Biological Sciences, Chinese Academy of Sciences, Shanghai 200031, China; CAS Key Laboratory of Systems Biology, CAS Center for Excellence in Molecular Cell Science, Institute of Biochemistry and Cell Biology, Shanghai Institutes for Biological Sciences, Chinese Academy of Sciences, University of Chinese Academy of Sciences, Shanghai, 200031, China; Bio-Med Big Data Center, Key Laboratory of Computational Biology, CAS-MPG Partner Institute for Computational Biology, Shanghai Institute of Nutrition and Health, Shanghai Institutes for Biological Sciences, University of Chinese Academy of Sciences, Chinese Academy of Sciences Shanghai, 200031, China; Center for Biomedical Informatics, Shanghai Children’s Hospital, Shanghai Jiao Tong University, Shanghai 200040, China; Department of Life Sciences, Shanghai Tech University, 100 Haike Road, Shanghai, 200031, China; School of Life Sciences and Biotechnology, Shanghai Jiao Tong University, Shanghai, 200240, China; Collaborative Innovation Center for Genetics and Development, Fudan University, Shanghai, China; Shanghai Center for Bioinformation Technology, Shanghai Academy of Science & Technology, Shanghai, China

## Abstract

Base editors hold promise for correcting pathogenic mutations, while substantial single nucleotide variations (SNVs) on both DNA and RNA were generated by cytosine base editors (CBEs). Here we examined possibilities to reduce off-target effects by engineering cytosine deaminases. By screening 24 CBEs harboring various rAPOBEC1 (BE3) or human APOBEC3A (BE3-hA3A) mutations on the ssDNA or RNA binding domain, we found 8 CBE variations could maintain high on-target editing efficiency. Using Genome-wide Off-target analysis by Two-cell embryo Injection (GOTI) method and RNA sequencing analysis, we found DNA off-target SNVs induced by BE3 could be completely eliminated in BE3^R126E^ but the off-target RNA SNVs was only slightly reduced. By contrast, BE3-hA3A^Y130F^ abolished the RNA off-target effects while could not reduce the DNA off-target effects. Notably, BE3^R132E^, BE3^W90Y+R126E^ and BE3^W90F+R126E^ achieved the elimination of off-target SNVs on both DNA and RNA, suggesting the feasibility of engineering base editors for high fidelity deaminases.

---

Base editors have been widely applied to perform targeted base editing and hold great potential for correcting pathogenetic mutations^1^. However, previous studies have identified off-target DNA edits by cytosine base editors (CBEs)^2, 3^, the most widely used cytosine base editors with rat APOBEC1 (rAPOBEC1) enzyme^4, 5^. Recently, several groups reported that CBEs with rAPOBEC1 (BE3) or human APOBEC3A (BE3-hA3A) can cause extensive transcriptome-wide RNA off-target edits in human cells^6–8^. These off-target RNA SNVs could be substantially decreased by screening CBEs harboring various rAPOBEC1 or hA3A mutations, but the DNA off-target edits of these variants were unknown^6–8^.

The observation of unwanted DNA and RNA off-target effects both have important implications for research and therapeutic applications of these technologies. Previous studies only examined the DNA (Zuo et al., 2019) or RNA off-target effects (Zhou et al., 2019) of base editors, here we analyzed both the DNA and RNA off-target effects of multiple engineered CBE variants by Genome-wide Off-target analysis by Two-cell embryo Injection (GOTI) and RNA-Seq analysis. We found that some variants could eliminate the DNA off-target activity while sustained RNA off-target effects. Conversely, some variant abolished the RNA off-target effects while maintained the DNA off-targets. Importantly, we successfully obtained three variants with the elimination of both DNA and RNA off-target effects.

We introduced various point mutations into rAPOBEC1 affecting the DNA^9–14^ or RNA^14, 15^ editing activity suggested by previous studies (Fig. 1a). The variants included deletions and mutations at the L-enriched 5’ or 3’ terminals of APOBEC1 (Del32, R33A, K34A, Del34, Del77, Del116, Del169, Del182, P190A and P191A), point mutations on the putative catalytic active site of APOBEC1 (H61A, H61R, V62A, E63A, E63Q, C93S, C96S). Based on the structure of human APOBEC3G^10, 11^, R126 is predicted to have interaction with the phosphate backbone of ssDNA (corresponding to R320 in APOBEC3G) (Fig. 1b, c). Compared with other mutations, R126E maintained on-target editing activity^9^. R128 and R132^9^ are near to R126 and could also affect the accessibility of ssDNA, so we also introduced mutations of R128E and R132E (Fig. 1a-c). We also examined the effect of combination of point mutations in the domain responsible for the hydrophobicity of the active site on APOBEC1 (W90A, W90F, W90Y), which was reported to narrow the width of base-editing window^9, 10^.

**Figure 1.**
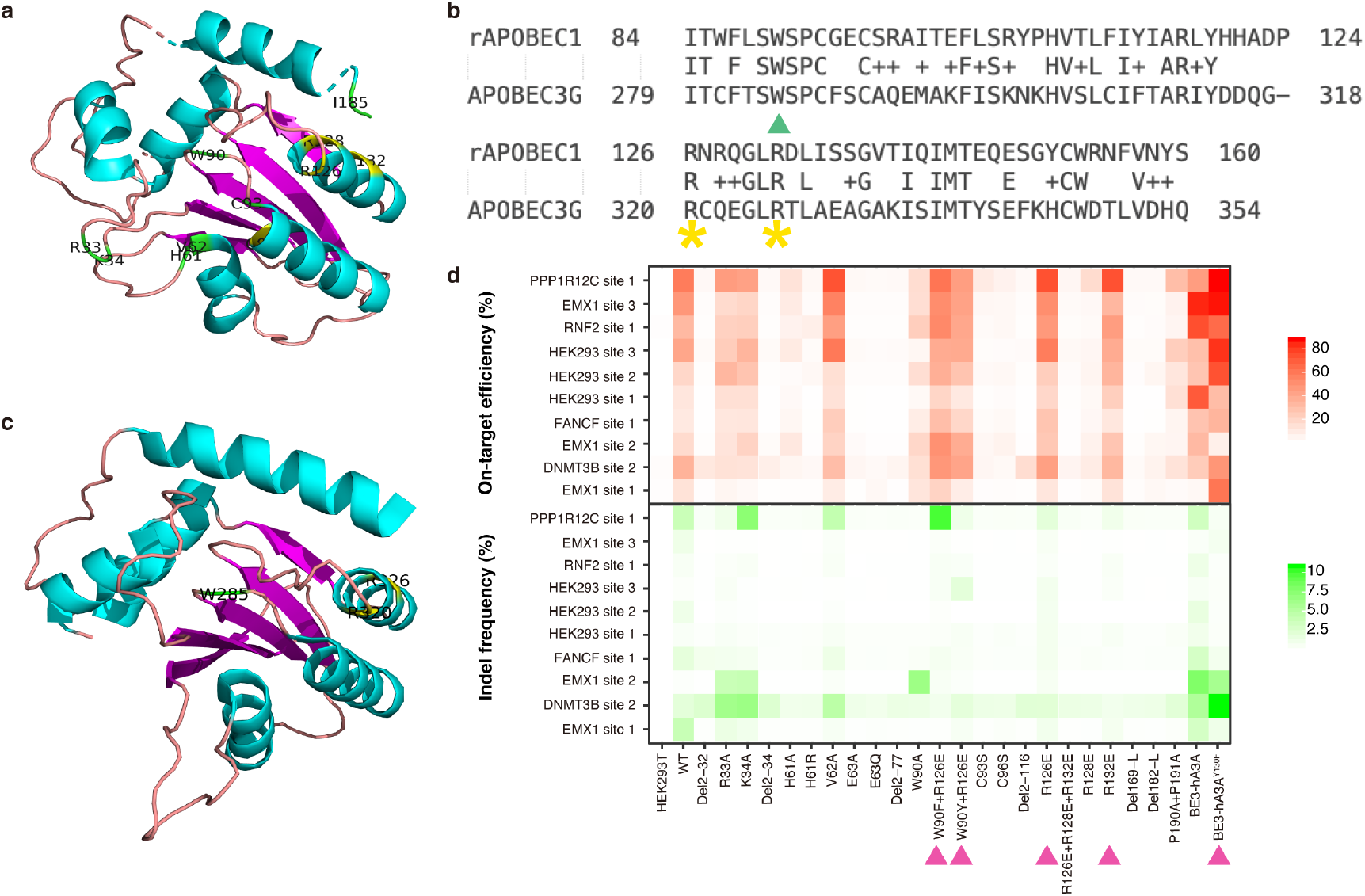
On-target efficiency of engineered CBEs. **a** The predicted structure of APOBEC1 with various rAPOBEC1 mutations. Mutated residues were highlighted and marked on the structure. **b** The sequence alignment between APOBEC3G and APOBEC1. Amino acid, identical residues; +, conservative substitutions. Green triangle represents residues in the hydrophobic active domain of APOBEC3G, and yellow stars indicate residues on the ssDNA binding domain. **c** The crystal structure of APOBEC3G. **d** The on-target efficiency and indel frequencies of different versions of engineered CBEs. Purple triangles indicate variants selected for the off-target evaluation. *n* = 3 biological replicates for each group.

We transfected HEK293T cells with plasmids encoding BE3 base editors harboring various mutations and evaluated their effects on both on-target efficiency and off-target rate. We tested the on-target activity of these variants on 10 genomic loci. Totally, by screening 23 engineered BE3 variants, we found 7 variants (R33A, K34A, V62A, W90F+R126E, W90Y+R126E, R126E and R132E) remained the on-target efficiency, and 4 of them (W90F+R126E, W90Y+R126E, R126E, R132E) showed no increase of indel rates (Fig. 1d and Supplementary Table 1). Besides, all of them showed no significantly difference on the editing window widths (Supplementary Fig. 1). Alternatively, we also tested one variant on hA3A (BE3-hA3A^Y130F^), reported to have high DNA on-target efficiency^16, 17^, and found it remained high on-target editing activities (Fig. 1d).

We next performed GOTI to evaluate the DNA off-target edits of the variants with high DNA on-target efficiency (BE3^R126E^, BE3^R132E^, BE3^W90Y+R126E^, BE3^W90F+R126E^ and BE3-hA3A^Y130F^) (Supplementary Table 2). The embryonic development was not affected by these variants injection except for BE3-hA3A (Supplementary Fig. 2). The on-target efficiency of these variants were confirmed by whole-genome sequencing (Fig. 2a). Notably, the number of DNA off-target SNVs in the embryos treated with BE3^R126E^, BE3^R132E^, BE3^W90Y+R126E^ or BE3^W90F+R126E^ was significantly reduced from 283 +/− 32 in wild-type BE3-treated embryos to 28 +/− 6 for BE3^R126E^, 43 +/− 11 for BE3^R132E^, 12 +/− 3 for BE3^W90Y+R126E^ and 39 +/− 27 for BE3^W90F+R126E^, similar to that found in non-edited control embryos (14 SNVs on average) and close to that of spontaneous mutation (Fig. 2b, Supplementary Fig. 3 and Supplementary Table 3). Besides, we observed no mutation bias and no SNVs that overlapped with the predicted off-target sites (Fig. 2c and Supplementary Fig. 4), suggesting the absence of DNA off-target SNVs induced by these variants. However, the BE3-hA3A^Y130F^ variant still generated substantial DNA off-target SNVs (409 +/− 86) (Fig. 2b and 2c).

**Figure 2.**
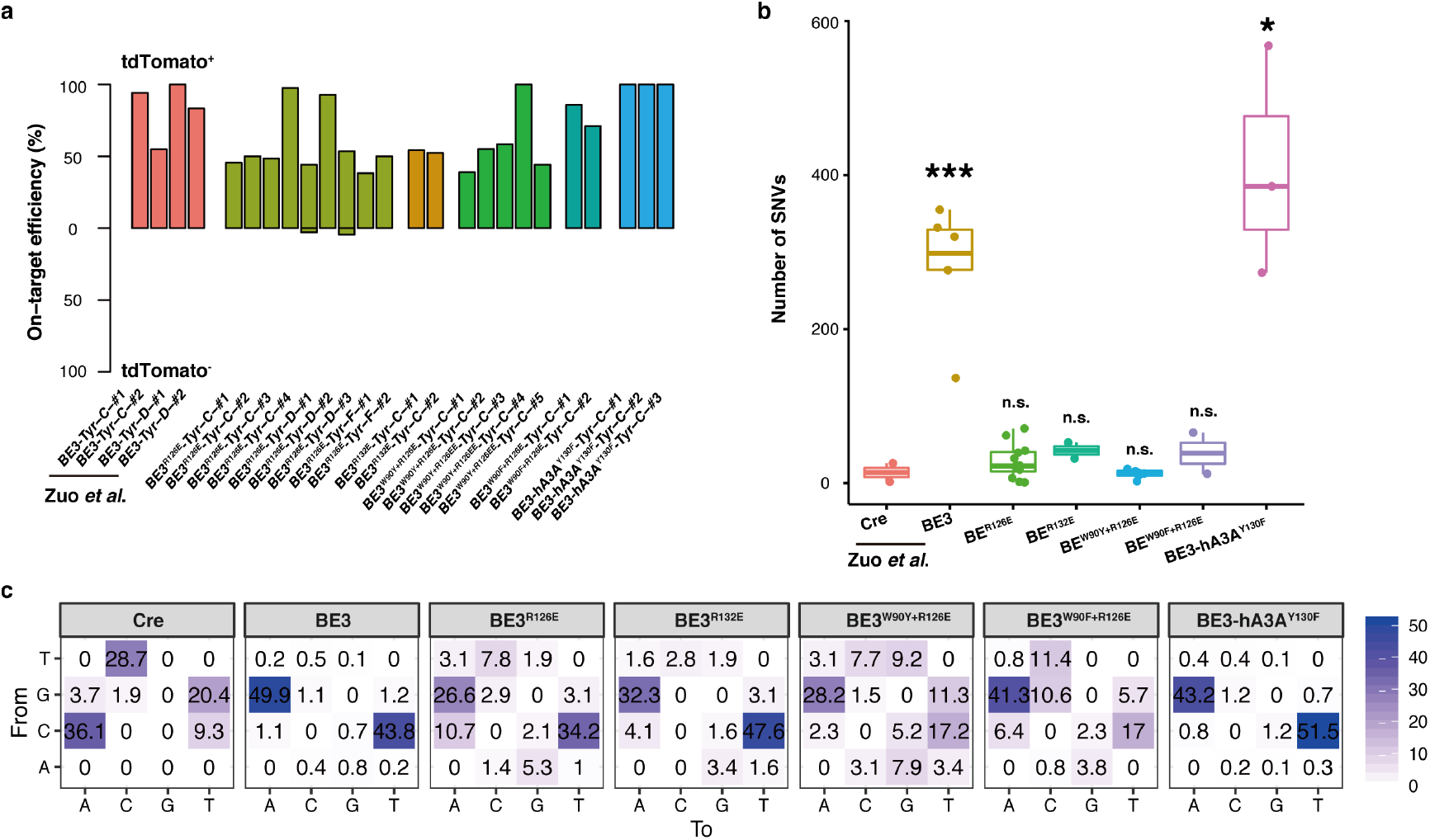
On-target and off-target evaluation of engineered CBEs by GOTI. **a** On-target efficiency of BE3 and CBE variants from WGS data. **b** The comparison of the total number of detected off-target SNVs. *n* = 2 for Cre, *n* = 6 for BE3, *n* = 10 for BE3^R126E^, *n* = 2 for BE3^R132E^, *n* = 5 for BE3^W90Y+R126E^, *n* = 2 for BE3^W90F+R126E^ and *n* = 3 for BE3-hA3A^Y130F^ groups. *P* value was calculated by two-sided Student’s t-test. \**P* < 0.05, \*\**P* < 0.01, \*\*\**P*<0.001. **c** Proportion of C>T and G>A mutations for Cre, BE3, and CBE variants-treated groups.

Moreover, we also evaluated the potential off-target effects on transcriptome of these variants. We found evident decrease of RNA off-target SNVs in BE3^R126E^, but the number was still significantly higher than that of the control group transfected with GFP (Fig. 3a and 3b). Intriguingly, two variants BE3^R132E^ and BE3^W90F+R126E^ (also know as BE3-FE1) ^9^ showed complete elimination of the RNA off-target edits. Combined with our previous results that BE3^W90Y+R126E^ (also know as BE3-YE1) ^9^ could completely eliminated the RNA off-target edits (Fig. 3a and 3b), we here obtained three variants, BE3^R132E^, BE3^W90Y+R126E^ and BE3^W90F+R126E^, with complete abolish of both DNA and RNA off-target effects.

**Figure 3.**
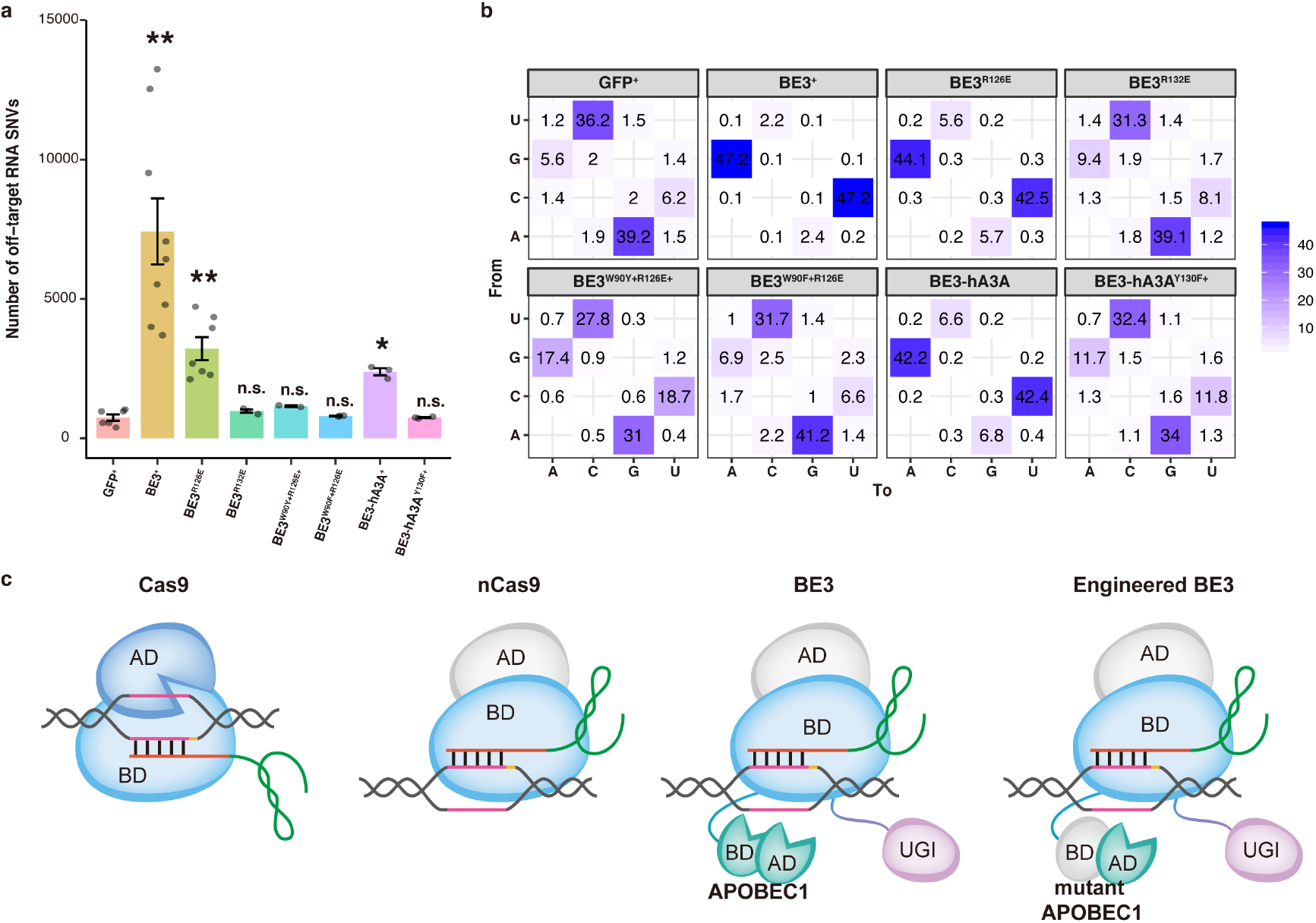
RNA off-target evaluation of engineered CBEs. **a** The comparison of the total number of detected RNA off-target SNVs. *n* = 6 for GFP, *n* = 8 for BE3, *n* = 6 for BE3^R126E^, *n* = 3 for BE3^R132E^, *n* = 2 for BE3^W90Y+R126E^, *n* = 3 for BE3^W90F+R126E^, *n* = 3 for BE3 (hA3A) and *n* = 3 for BE3-hA3A^Y130F^ groups. *P* value was calculated by two-sided Student’s t-test. \**P* < 0.05, \*\**P* < 0.01, \*\*\**P*<0.001. **b** Proportion of C>U and G>A mutations for GFP, BE3, and BE3 variants-treated groups. **c** Model of CBE optimization. The nickase Cas9 (nCas9) of engineered CBE loses one nuclease activity of Cas9 while remains the DNA binding ability. In contrast to nCas9, mutant APOBEC1 of engineered CBE loses the binding ability of ssDNA and RNA but remains the deaminase activity. AD, active domain; BD, binding domain; APOBEC1, rAPOBEC1; UGI, uracil DNA glycosylase inhibitor.

Considering that GOTI was developed to examine the sgRNA-independent off-target effects, we also examined the sgRNA-dependent off-target sites as previous studies^18^. We found no increase of the number of these sgRNA-dependent off-targets in all the variants (Supplementary Fig. 5).

In summary, by screening dozens of mutations on rAPOBEC1 or hA3A from multiple researches before, we found three variants with complete abolish of both DNA and RNA off-targets with no compromise for on-target activity. Although BE3^R132E^, BE3^W90Y+R126E^ and BE3^W90F+R126E^ have been reported to remain editing efficiencies as BE3^9^, off-target evaluation is necessary for their clinical application. In addition, we found that BE3^R126E^ could eliminate the DNA off-target effects but not the RNA off-targets, while BE3-hA3A^Y130F^ only reduced the RNA off-target effects, indicating that the elimination of DNA off-target effects was not eligible for the minimization of RNA off-target effects, and vice versa. Engineered variants with high fidelity on both DNA and RNA provide a safe tool for gene editing. Notably, the study described here demonstrates that the DNA and RNA off-target effects of BE3 could be simultaneously eliminated by engineering APOBEC1 with mutations on the putative ssDNA binding domain and hydrophobic domain but not on catalytic domain. Therefore, our work illustrates how the off-target effects can be defined and minimized for research and therapeutic applications (Fig. 3c). This approach for fusion protein optimization could be generalized in other synthetic tools such as CRISPR/Cas9 derivates (Supplementary Fig. 6).

## Acknowledgments

We thank FACS facility HW, LQ, SQ in ION and MZ in IPS, LY in Big Data Platform (SIBS,CAS). These work were supported by R&D Program of China (2018YFC2000100 and 2017YFC1001302), CAS Strategic Priority Research Program (XDB32060000), National Natural Science Foundation of China (31871502, 31522037, 31822035), Shanghai Municipal Science and Technology Major Project (2018SHZDZX05), Shanghai City Committee of science and technology project (18411953700, 18JC1410100), National Science and Technology Major Project (2015ZX10004801-005), National Key Research and Development Program of China (2017YFA0505500, 2016YFC0901704) and the Agricultural Science and Technology Innovation Program.

## Author contributions

EZ designed and performed experiments. YS, WW, RZ and LY performed data analysis. TY, BH and JL performed PCR analysis. WY performed mouse embryo transfer. HY and YL supervised the project and designed experiments. YS and HY wrote the paper.

## Data and materials availability

All the sequencing data were deposited in NCBI Sequence Read Archive (SRA) under project accession PRJNA527003.

## Supplementary Information

### Materials and Methods

#### Animal care

Heterozygous Ai9 (full name B6.Cg-Gt (ROSA) 26Sortm9 (CAG-td-Tomato) Hze/J; JAX strain 007909) male mice and female C57BL/6 mice (4 weeks old) were mated for embryo collection. ICR females were used for recipients. The animals usage and care complied with the guideline of the Biomedical Research Ethics Committee of Shanghai Institutes for Biological Science, Chinese Academy of Sciences.

#### Generation of mutant base editor mRNA and sgRNA

T7 promoter was added to base editor coding region by PCR amplification of plasmid, using primer base editor F and R. T7-base editor PCR product was purified and used as the template for in vitro transcription (IVT) using mMESSAGE mMACHINE T7 ULTRA kit (Life Technologies). T7 promoter was added to sgRNA template by PCR amplification of px330. The T7-sgRNA PCR product was purified and used as the template for IVT using MEGA shortscript T7 kit (Life Technologies). T7 promoter was added to Cre template by PCR amplification. T7-Cre PCR product was purified and used as the template for *in vitro* transcription (IVT) using mMESSAGE mMACHINE T7 ULTRA kit (Life Technologies). Cas9 mRNA, Cre mRNA and sgRNAs were purified using MEGA clear kit (Life Technologies) and eluted in RNase-free water. sgRNA sequences

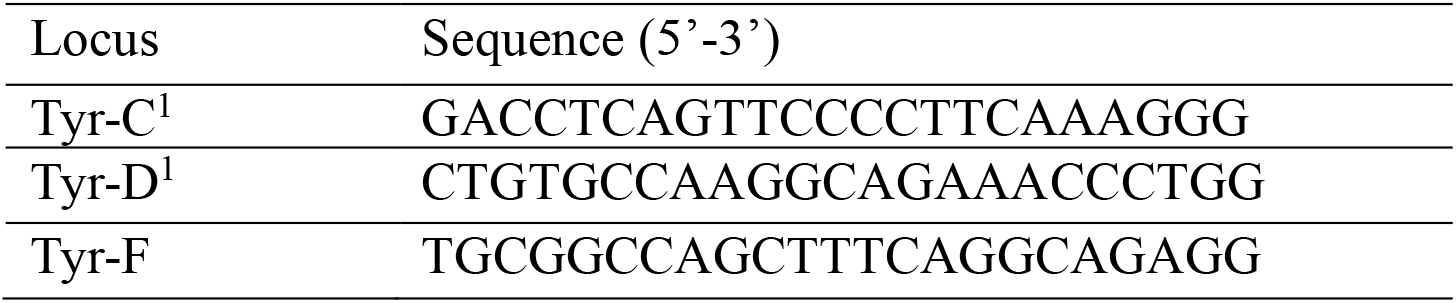

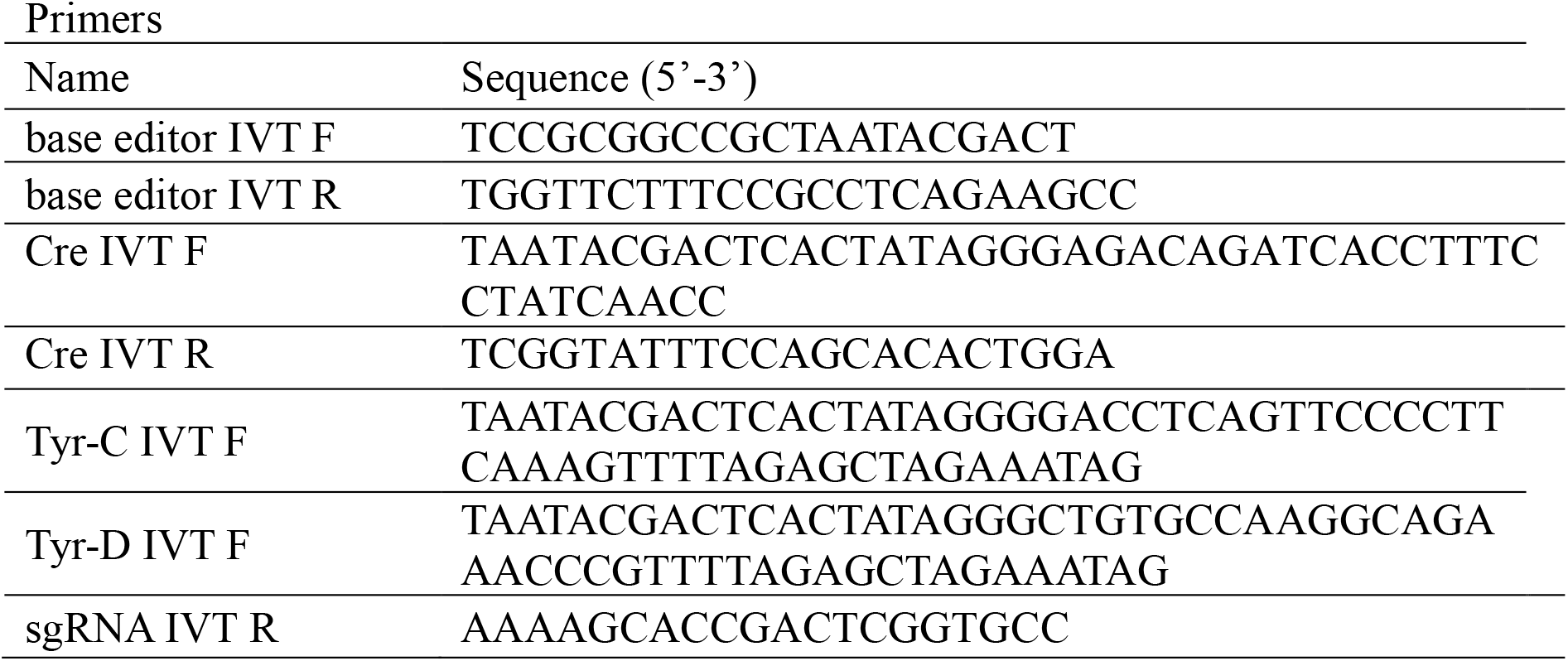

#### 2-cell Embryo Injection, Embryo Culturing, and Embryo Transplantation

Super ovulated C57BL/6 females (4 weeks old) were mated to heterozygous Ai9 (full name B6.Cg-Gt(ROSA)26Sortm9(CAG-td-Tomato)Hze/J; JAX strain 007909) males, and fertilized embryos were collected from oviducts 23 h post hCG injection. For 2-cell editing, the mixture of BE3 mRNA (10 or 50 ng/μl) or BE3^R126E^ mRNA (50 ng/μl), sgRNA (50 ng/μl) and Cre mRNA (2 ng/μl) was injected into the cytoplasm of one blastomere of 2-cell embryo 48 h post hCG injection in a droplet of M2 medium containing 5 μg/ml cytochalasin B (CB) using a FemtoJet microinjector (Eppendorf) with constant flow settings. The injected embryos cultured in KSOM medium with amino acids at 37 °C under 5% CO_2_ in air for 2 hours and then transferred into oviducts of pseudopregnant ICR females at 0.5 dpc.

#### Cloning

Site-directed mutagenesis of BE3 was done using NEBuilder HiFi DNA Assembly Master Mix (New England BioLabs). Briefly, a primer with an overhang containing the desired point mutation was used to amplify the appropriate vector plasmid by PCR. pCMV-BE3 variants-polyA-pCMV-mCherry-polyA was generated through NEBuilder HiFi DNA Assembly, by combining a PCR-amplified pCMV-mCherry-poly A with a digested pCMV-BE3 variants backbone. pCMV-EGFP-polyA-U6-sgRNA were generated through NEBuilder HiFi DNA Assembly, by combining a PCR-amplified U6-sgRNA with a digested pCMV-EGFP-poly A backbone.

#### Cell culture, transfections and FACS

HEK293T cells were maintained in Dulbecco’s modified eagle medium (DMEM) supplemented with 10% fetal bovine serum (FBS) in a 37°C humidified incubator with 5% CO2. pCMV-BE3 (WT/BE3 variants)-polyA-pCMV-mCherry-polyA and pCMV-EGFP-polyA-U6-sgRNA expression plasmids were co-transfected using Lipofectamine 3000 (ThermoFisher Scientific) according to the manufacturer’s protocol. 72 hr post transfection, cells were washed with phosphate buffered saline (PBS) and trypsinized using 0.05% trypsin-EDTA. Cell suspension was filtered through a 40-μm cell strainer, and EGFP/mCherry positive cells were isolated by FACS.

#### FACS

To isolate mouse embryonic cells, the prepared tissues were dissociated enzymatically in an incubation solution of 5 mL Trypsin-EDTA (0.05%) at 37°C for 30min. The digestion was stopped by adding 5 ml of DMEM medium with 10% Fetal Bovine Serum (FBS). Fetal tissues were then homogenized by passing 30-40 times through a 1ml pipette tips. The cell suspension was centrifuged for 6 min (800 rpm), and the pellet was resuspended in DMEM medium with 10% FBS. Finally, the cell suspension was filtered through a 40μm cell strainer, and tdtomato^+^/tdtomato^−^ cells were isolated by FACS. Samples were found to be >95% pure when assessed with a second round of flow cytometry and fluorescence microscopy analysis.

#### Whole genome sequencing and data analysis

DNeasy blood and tissue kit (catalog number 69504, Qiagen) was used to extract genomic DNA from cells following the manufacturer’s instructions. Whole genome sequencing was performed at mean coverages of 50x by Illumina HiSeq X Ten. BWA (v0.7.12) was used to map qualified sequencing reads to the reference genome (mm10). The mapped BAM files were then sorted and marked using Picard tools (v2.3.0). To identify the genome wide *de novo* SNVs with high confidence, we conducted single nucleotide variation calling on three algorithms, Mutect2 (v3.5), Lofreq (v2.1.2) and Strelka (v2.7.1), separately ^2–4^. In parallel, Mutect2 (v3.5), Scalpel (v0.5.3) and Strelka (v2.7.1) were run individually for the detection of whole genome *de novo* indels ^2, 4, 5^. The overlap of three algorithms of SNVs or indels were considered as the true variants. All the sequencing data were deposited in NCBI Sequence Read Archive (SRA) under project accession PRJNA527003.

Potential off-targets of targeted sites were predicted using two previous reported algorithms, Cas-OFFinder (http://www.rgenome.net/cas-offinder/) and CRISPOR (http://crispor.tefor.net/) with all possible mismatches ^6, 7^.

The SNVs and indels were annotated with annovar (version 2016-02-01) using RefSeq database ^8^.

#### Structure prediction

Amino acid sequences of rat APOBEC1 and human APOBEC3G were retrieved from UniProt (https://www.uniprot.org/) and sequence alignment was performed with NCBI blastp (https://blast.ncbi.nlm.nih.gov/Blast.cgi?PROGRAM=blastp&PAGE_TYPE=BlastSearch&LINK_LOC=blasthome). The structure of rAPOBEC1 was predicted by protein structure prediction server, (PS)^2 9, 10^ according to the consensus sequence and secondary structure information for proteins with known structures. The crystal structure of APOBEC3G was downloaded from PDB (http://www.rcsb.org/3d-view/3IQS) and presented using PyMOL (v2.3.2).

#### Statistical analysis

R version 3.5.1 (http://www.R-project.org/) was used to conduct all the statistical analyses in this work. All tests conducted were two-sided, and the significant difference was considered at *P* < 0.05.

---

**Supplementary Figure 1.**
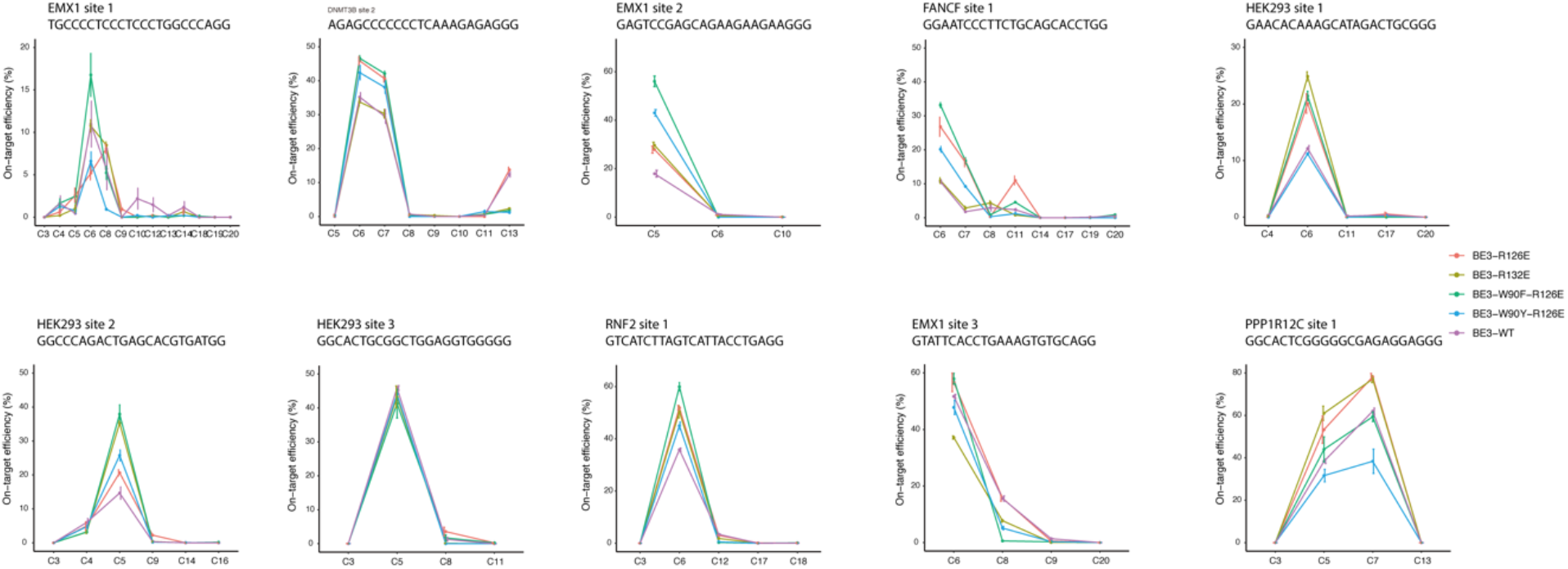
The editing window of the BE3 and BE3 variants in different target sites. *n* = 3 biological replicates for each group.

**Supplementary Figure 2.**
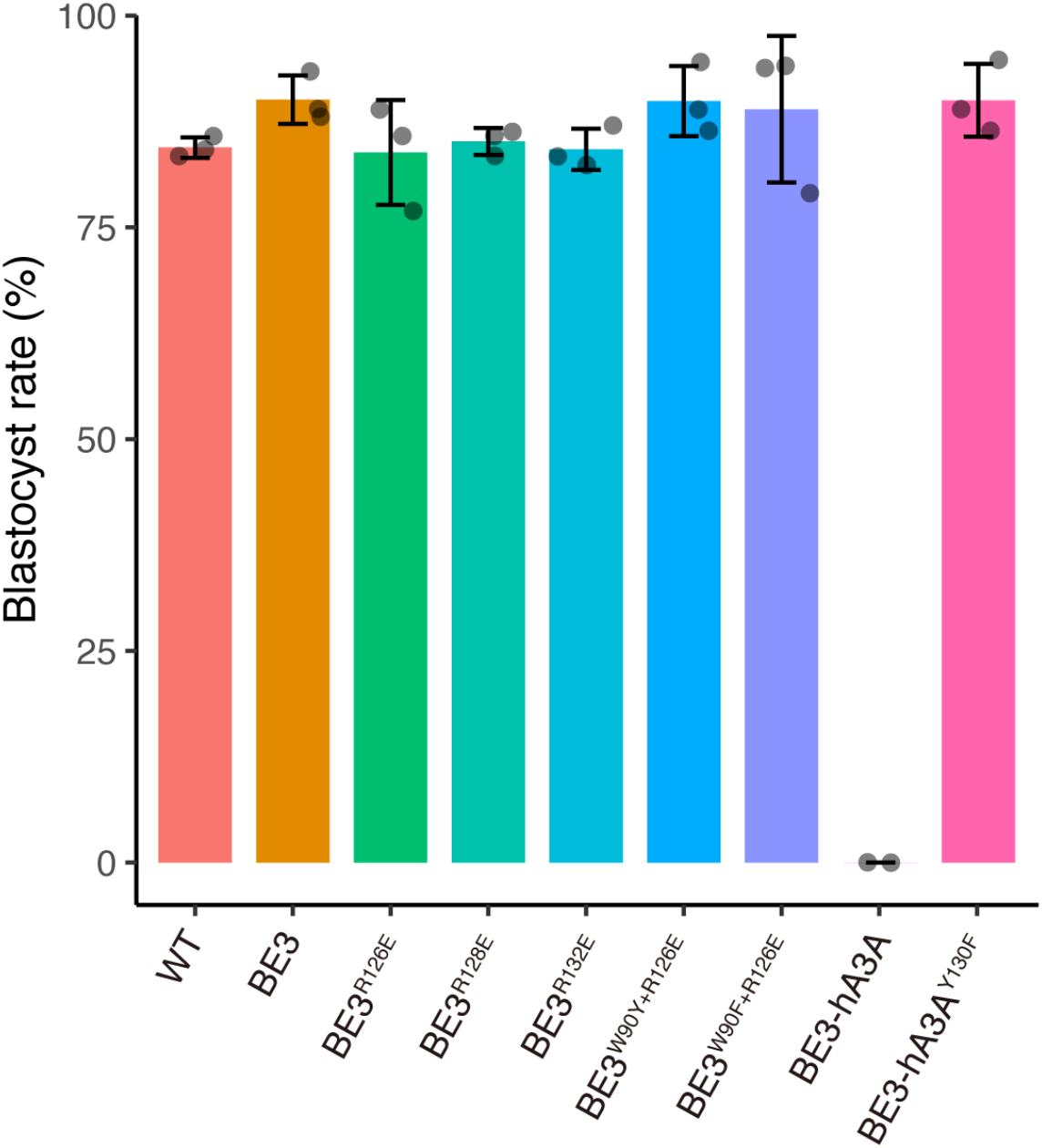
The embryonic development rates for BE3 and BE3 variants. *n* = 3 biological replicates for each group.

**Supplementary Figure 3.**
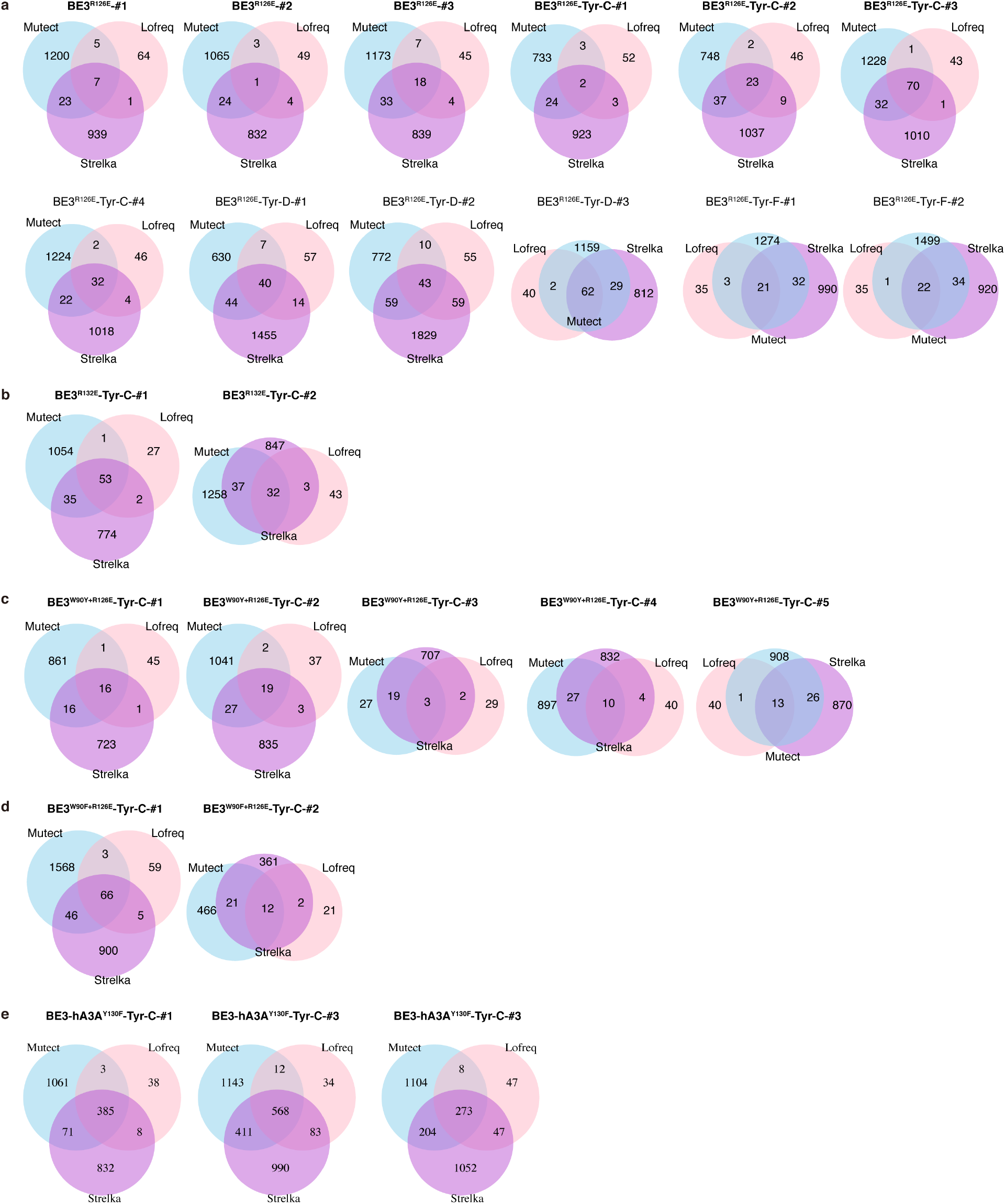
Venn diagrams of SNVs detected in each embryo by WGS data using the indicated software tools. **a** SNVs identified in BE3^R126E^-treated embryos. **b** SNVs identified in BE3^R132E^-treated embryos. **c** SNVs identified in BE3^W90Y+R126E^-treated embryos. **d** SNVs identified in BE3^W90F+R126E^-treated embryos. **e** SNVs identified in BE3-hA3A^Y130F^-treated embryos.

**Supplementary Figure 4.**
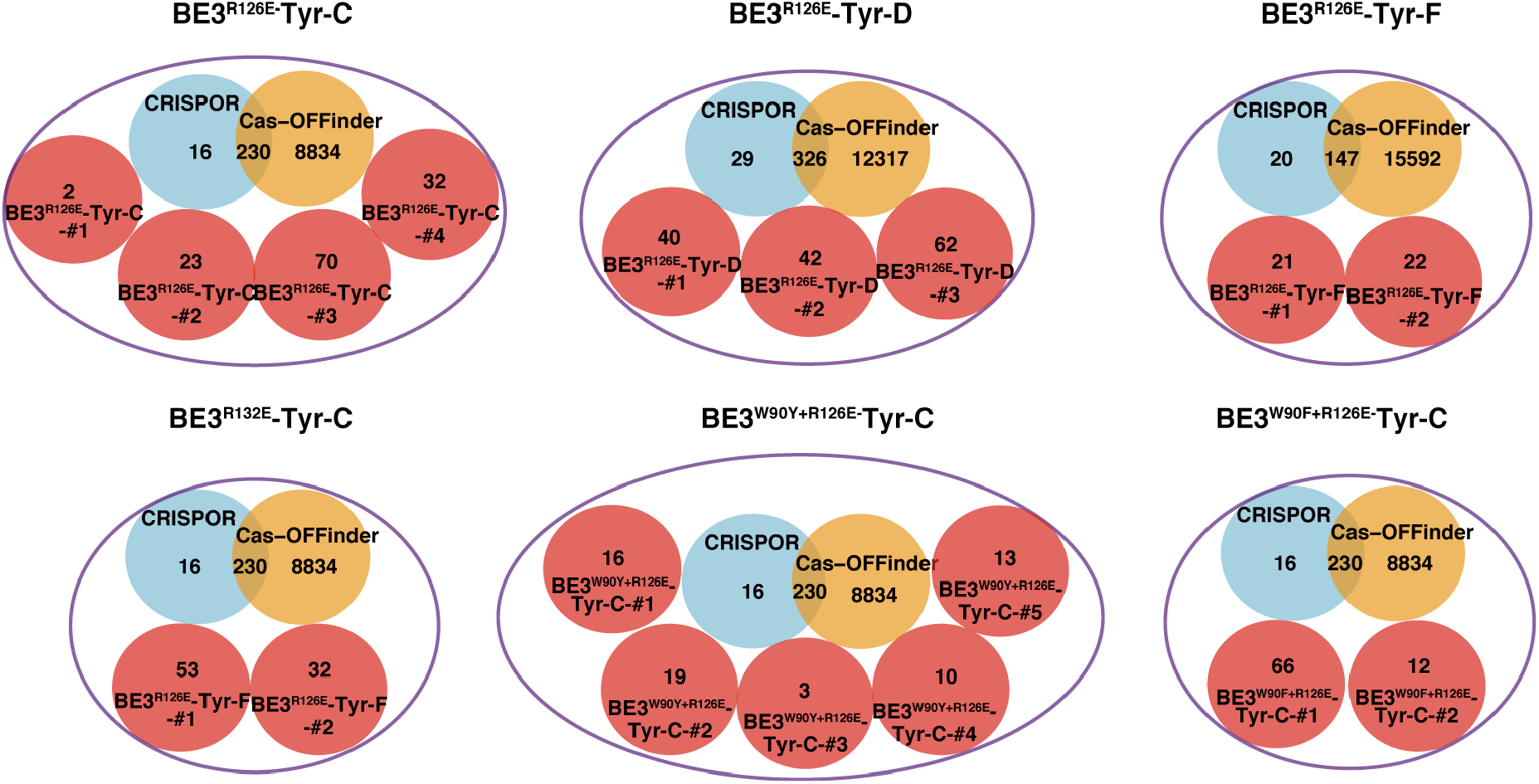
The overlap among SNVs detected from our analysis with predicted off-targets sites by Cas-OFFinder and CRISPOR.

**Supplementary Figure 5.**
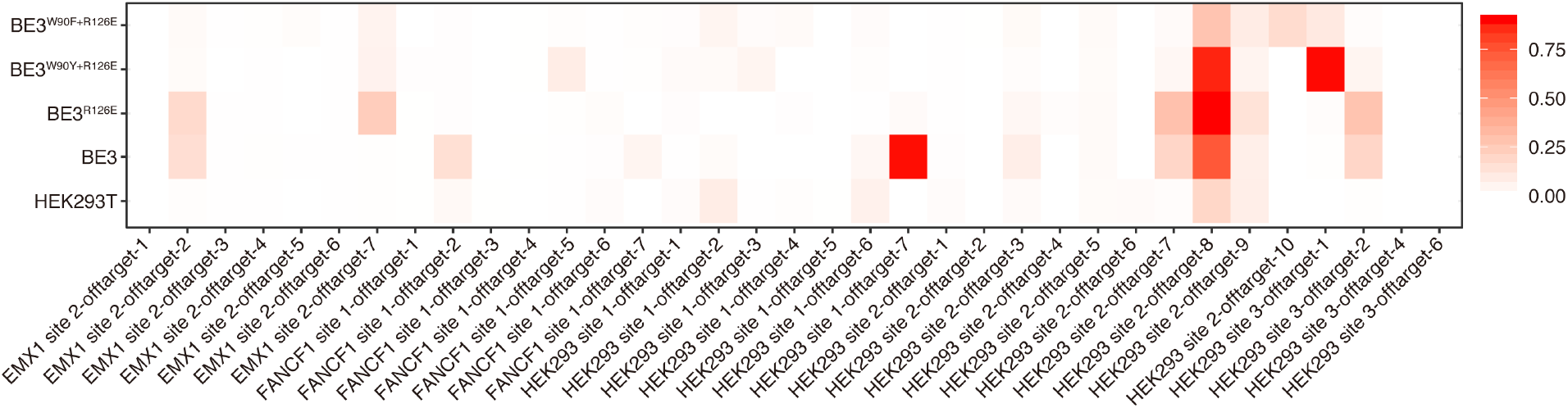
Activities of BE3 and BE3 variants at the indicated off-target sites. HEK293T cells were transfected with plasmids expressing BE3, BE3^R126E^, BE3^W90Y+R126E^ or BE3 ^W90F+R126E^ and sgRNAs matching the indicated on-target sequence using Lipofectamine 3000. Three days after transfection, genomic DNA was extracted, amplified by PCR, and analyzed by high-throughput DNA sequencing at the on-target loci, plus the top ten known Cas9 off-target loci for these sgRNAs, as previously determined using the GUIDE-seq method ^11, 12^ and ChIP-seq method ^13^. Sequences of the on-target and off-target protospacers and primers were shown in Table S5. Each cell represents the percentage of total sequencing reads with C to T conversion. *n* = 3 biological replicates for each group.

**Supplementary Figure 6.**
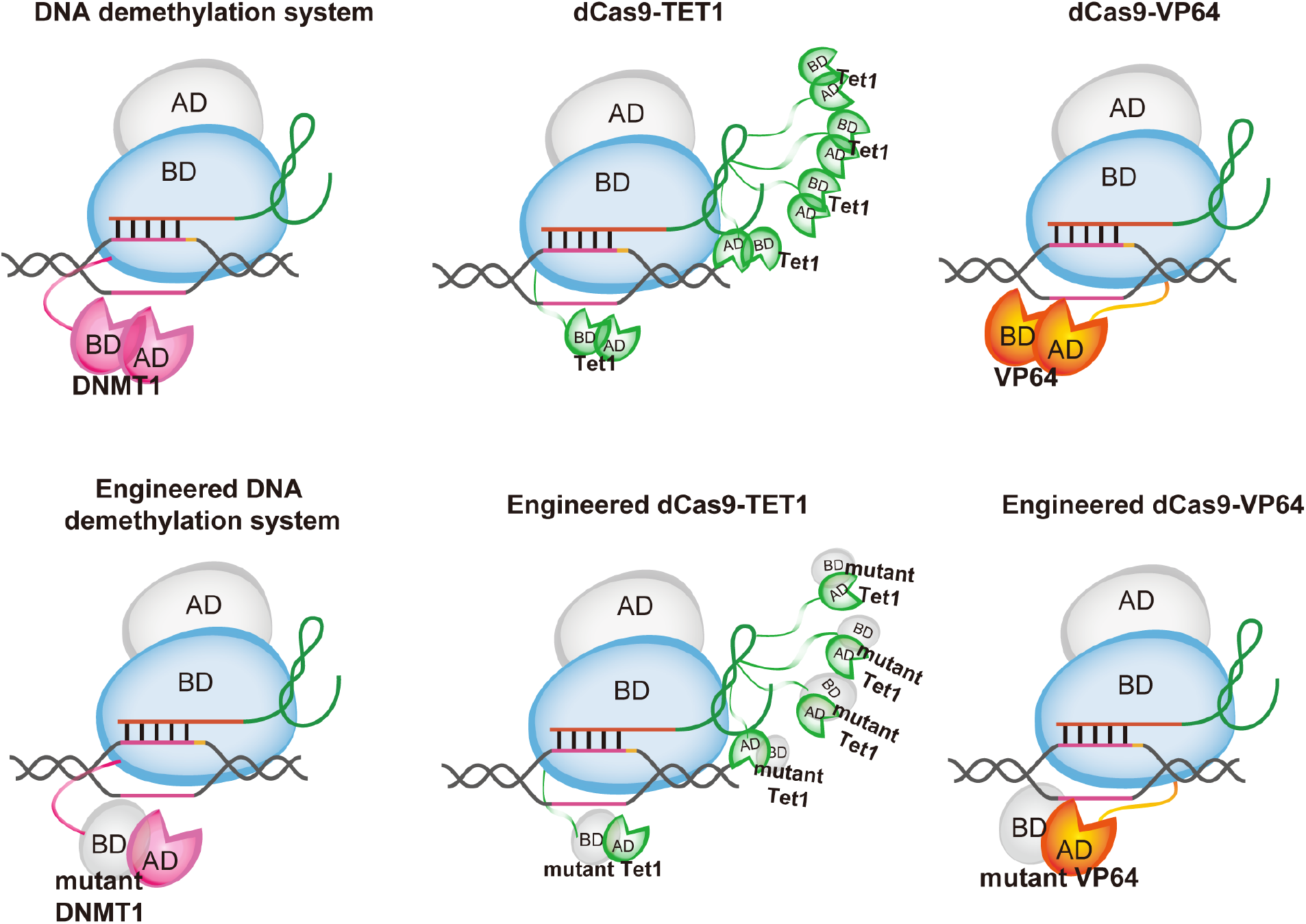
The generalization of optimization method for other CRISPR/Cas9 derivates. AD, active domain; BD, binding domain; Tet1, Ten-Eleven Translocation dioxygenase 1.

**Supplementary Table 1.** P values of the on-target efficiency and indel rates between CBE variants and BE3.

| **Mutant** | **On-target efficiency** | **Indel frequency** |
| --- | --- | --- |
| **Del2-32** | 0.001 | 0.046 |
| **R33A** | 0.191 | 0.957 |
| **K34A** | 0.257 | 0.605 |
| **R33A+K34A** | 0.748 | 0.020 |
| **Del2-34** | 0.001 | 0.032 |
| **H61A** | 0.002 | 0.014 |
| **H61R** | 0.001 | 0.015 |
| **V62A** | 0.517 | 0.724 |
| **E63A** | 0.001 | 0.015 |
| **E63Q** | 0.001 | 0.014 |
| **Del2-77** | 0.001 | 0.017 |
| **W90A** | 0.019 | 0.411 |
| **W90F+R126E** | 0.095 | 0.893 |
| **W90Y+R126E** | 0.717 | 0.126 |
| **C93S** | 0.001 | 0.016 |
| **C96S** | 0.001 | 0.018 |
| **Del2-116** | 0.001 | 0.026 |
| **R126E** | 0.270 | 0.282 |
| **R126E+R132E** | 0.430 | 0.027 |
| **R126E+R128E+R132E** | 0.001 | 0.018 |
| **R128E** | 0.002 | 0.016 |
| **R132E** | 0.563 | 0.036 |
| **Del169-L** | 0.001 | 0.015 |
| **Del-182-L** | 0.001 | 0.015 |
| **P190A+P191A** | 0.014 | 0.029 |
| **A3A** | 0.197 | 0.180 |
| **hA3A-Y130F** | 0.200 | 0.628 |

**Supplementary Table 2.** Summary of HiSeq X Ten sequencing.

| Sample | Sample Code | Group | Mapped bases (Gbp) | Coverage |
| --- | --- | --- | --- | --- |
| **BE3<sup>R126E</sup>-#1** | A21 | tdTomato+ | 120.21 | 43.38 |
|  | A22 | tdTomato- | 118.12 | 42.63 |
| **BE3<sup>R126E</sup>-#2** | A23 | tdTomato+ | 124.13 | 44.80 |
|  | A24 | tdTomato- | 121.36 | 43.80 |
| **BE3<sup>R126E</sup>-#3** | A185 | tdTomato+ | 115.12 | 41.55 |
|  | A186 | tdTomato- | 117.56 | 42.43 |
| **BE3<sup>R126E</sup>-Tyr-C-#1** | A29 | tdTomato+ | 141.77 | 51.16 |
|  | A30 | tdTomato- | 143.94 | 51.94 |
| **BE3<sup>R126E</sup>-Tyr-C-#2** | A35 | tdTomato+ | 146.12 | 52.73 |
|  | A36 | tdTomato- | 143.13 | 51.65 |
| **BE3<sup>R126E</sup>-Tyr-C-#3** | A225 | tdTomato+ | 141.81 | 51.18 |
|  | A226 | tdTomato- | 136.56 | 49.28 |
| **BE3<sup>R126E</sup>-Tyr-C-#4** | A227 | tdTomato+ | 134.54 | 48.55 |
|  | A228 | tdTomato- | 130.00 | 46.92 |
| **BE3<sup>R126E</sup>-Tyr-D-#1** | A123 | tdTomato+ | 113.09 | 40.81 |
|  | A124 | tdTomato- | 109.02 | 39.34 |
| **BE3<sup>R126E</sup>-Tyr-D-#2** | A131 | tdTomato+ | 107.61 | 38.83 |
|  | A132 | tdTomato- | 127.93 | 46.17 |
| **BE3<sup>R126E</sup>-Tyr-D-#3** | A141 | tdTomato+ | 118.05 | 42.60 |
|  | A142 | tdTomato- | 119.45 | 43.11 |
| **BE3<sup>R126E</sup>-Tyr-F-#1** | A251 | tdTomato+ | 146.49 | 52.87 |
|  | A252 | tdTomato- | 129.56 | 46.76 |
| **BE3<sup>R126E</sup>-Tyr-F-#2** | A258 | tdTomato+ | 142.19 | 51.31 |
|  | A259 | tdTomato- | 114.74 | 41.41 |
| **BE3<sup>W90Y+R126E</sup>-Tyr-C-#1** | A267 | tdTomato+ | 119.09 | 42.98 |
|  | A268 | tdTomato- | 142.76 | 51.52 |
| **BE3<sup>W90Y+R126E</sup>-Tyr-C-#2** | A269 | tdTomato+ | 142.76 | 51.52 |
|  | A270 | tdTomato- | 122.23 | 44.11 |
| **BE3<sup>W90Y+R126E</sup>-Tyr-C-#3** | A271 | tdTomato+ | 116.99 | 42.22 |
|  | A272 | tdTomato- | 137.70 | 49.69 |
| **BE3<sup>W90Y+R126E</sup>-Tyr-C-#4** | A273 | tdTomato+ | 126.17 | 45.53 |
|  | A274 | tdTomato- | 148.42 | 53.56 |
| **BE3<sup>W90Y+R126E</sup>-Tyr-C-#5** | A275 | tdTomato+ | 130.04 | 46.93 |
|  | A276 | tdTomato- | 146.61 | 52.91 |
| **BE3<sup>W90Y+R126E</sup>-Tyr-C-#1** | A301 | tdTomato+ | 130.76 | 47.19 |
|  | A302 | tdTomato- | 135.93 | 49.05 |
| **BE3<sup>W90Y+R126E</sup>-Tyr-C-#2** | A303 | tdTomato+ | 143.04 | 51.62 |
|  | A304 | tdTomato- | 149.73 | 54.04 |
| **BE3<sup>W90F+R126E</sup>-Tyr-C-#1** | A307 | tdTomato+ | 147.88 | 53.37 |
|  | A308 | tdTomato- | 117.40 | 42.37 |
| **BE3<sup>W90F+R126E</sup>-Tyr-C-#2** | A309 | tdTomato+ | 83.88 | 30.27 |
|  | A310 | tdTomato- | 126.48 | 45.64 |
| **BE3-hA3A<sup>Y130F</sup>-Tyr-C-#1** | A277 | tdTomato+ | 137.38 | 49.58 |
|  | A278 | tdTomato- | 152.07 | 54.88 |
| **BE3-hA3A<sup>Y130F</sup>-Tyr-C-#2** | A281 | tdTomato+ | 147.01 | 53.05 |
|  | A282 | tdTomato- | 149.36 | 53.90 |
| **BE3-hA3A<sup>Y130F</sup>-Tyr-C-#3** | A283 | tdTomato+ | 149.70 | 54.02 |
|  | A284 | tdTomato- | 151.43 | 54.65 |

**Supplementary Table 4.**
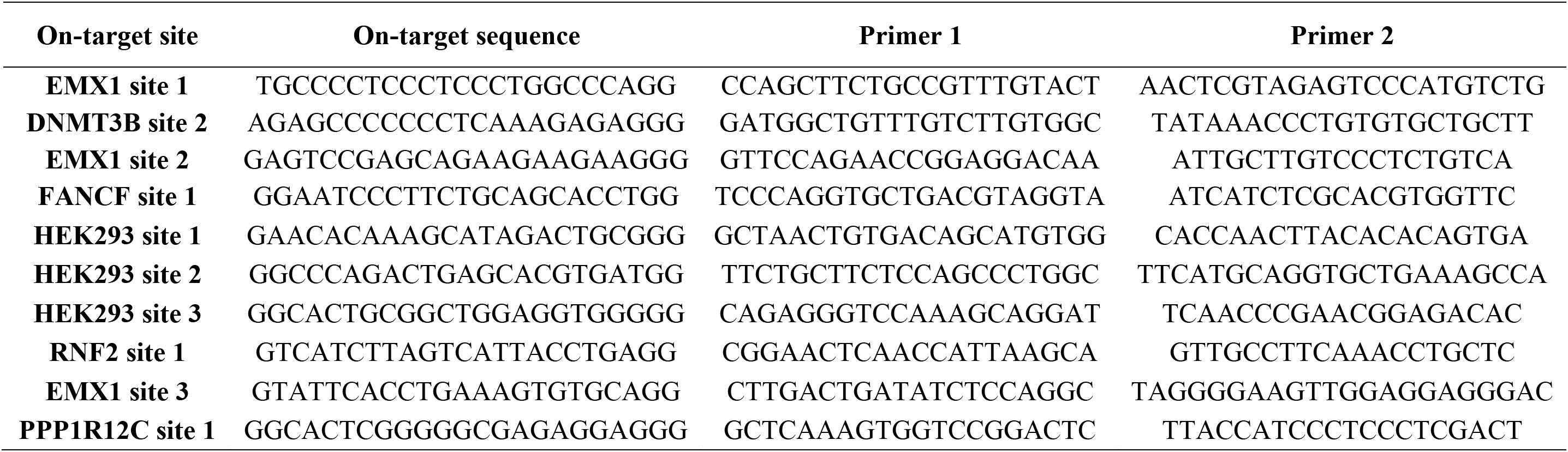
Primers used for deep sequencing of on-target activity.

**Supplementary Table 5.**
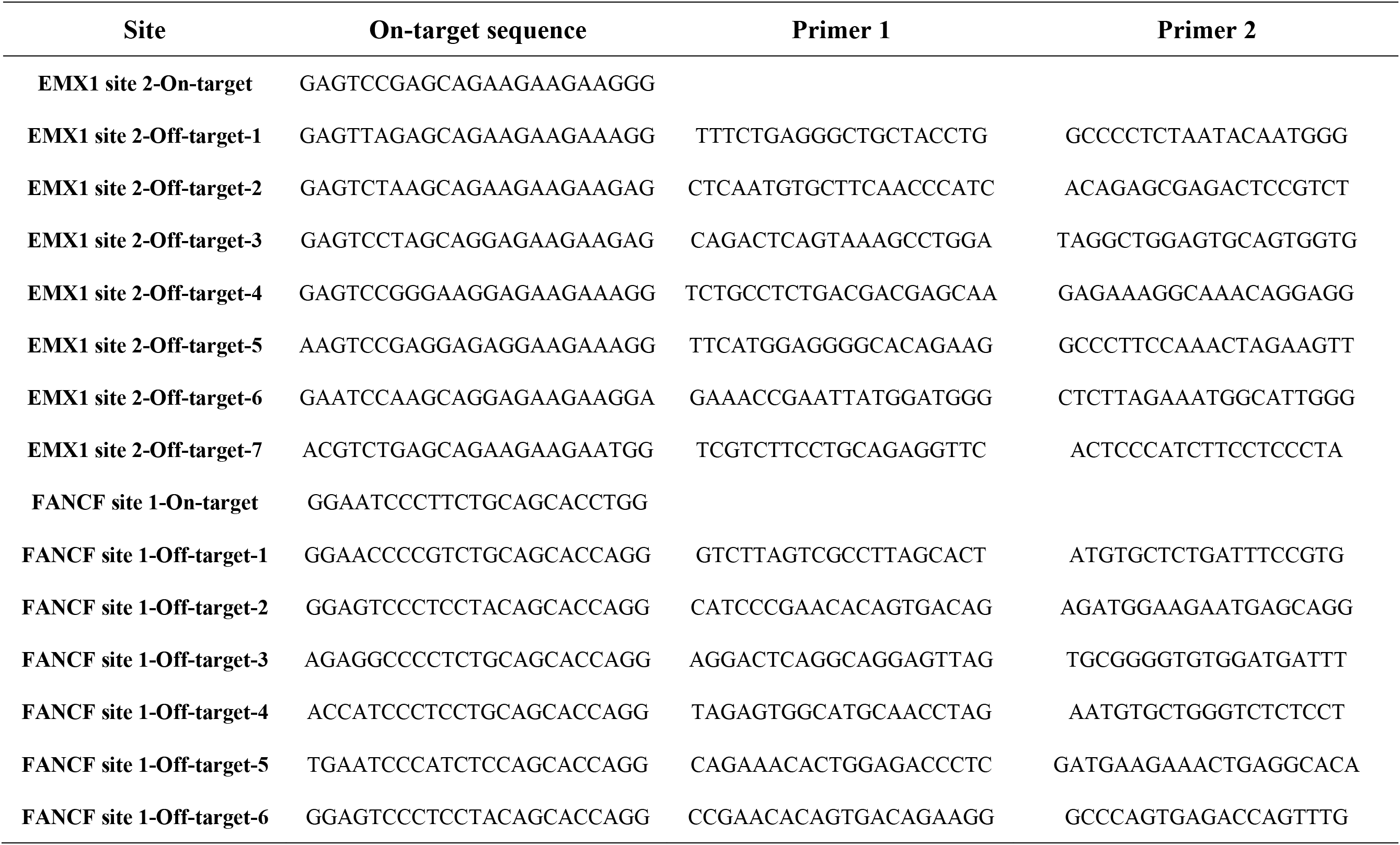
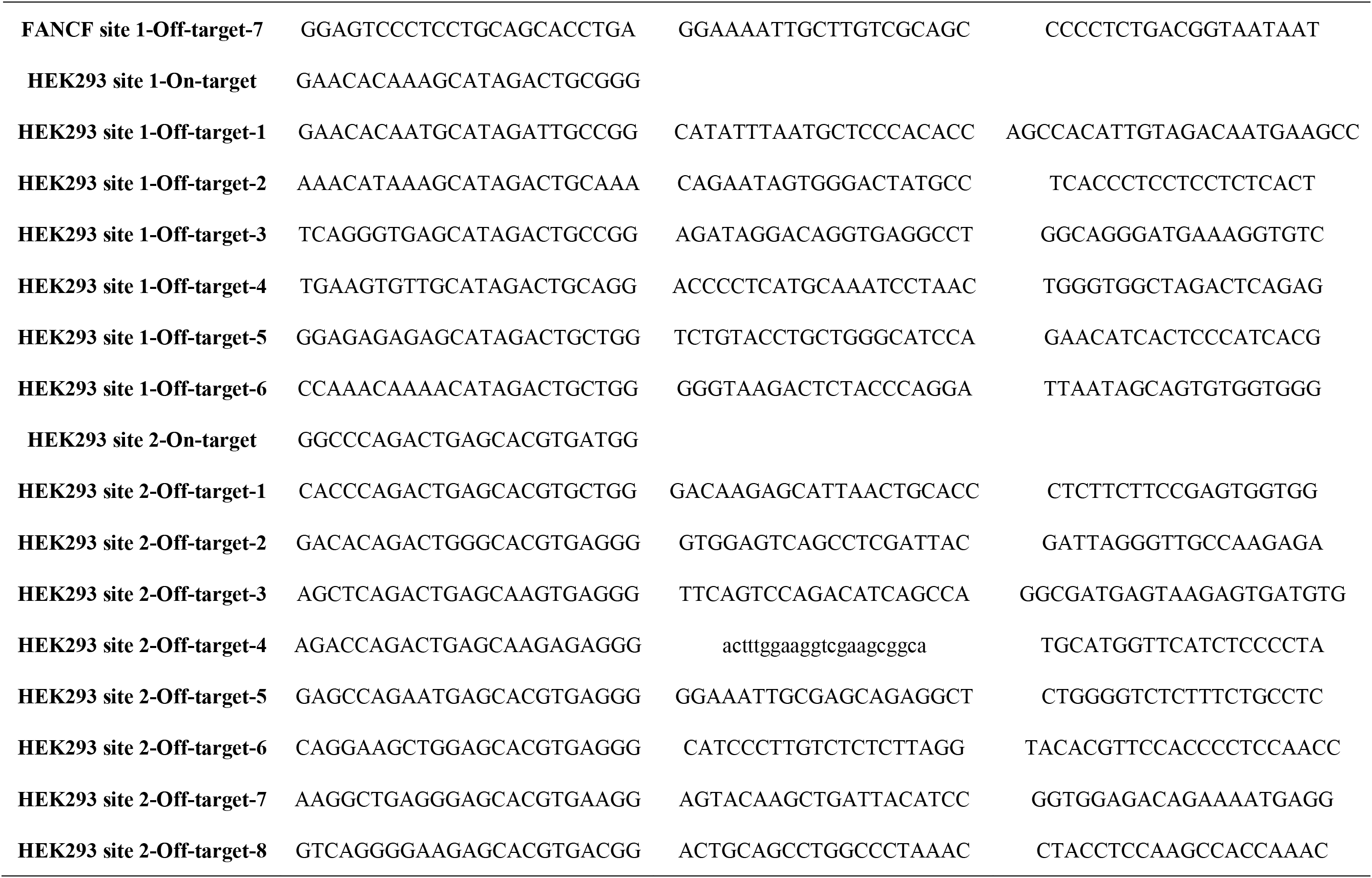
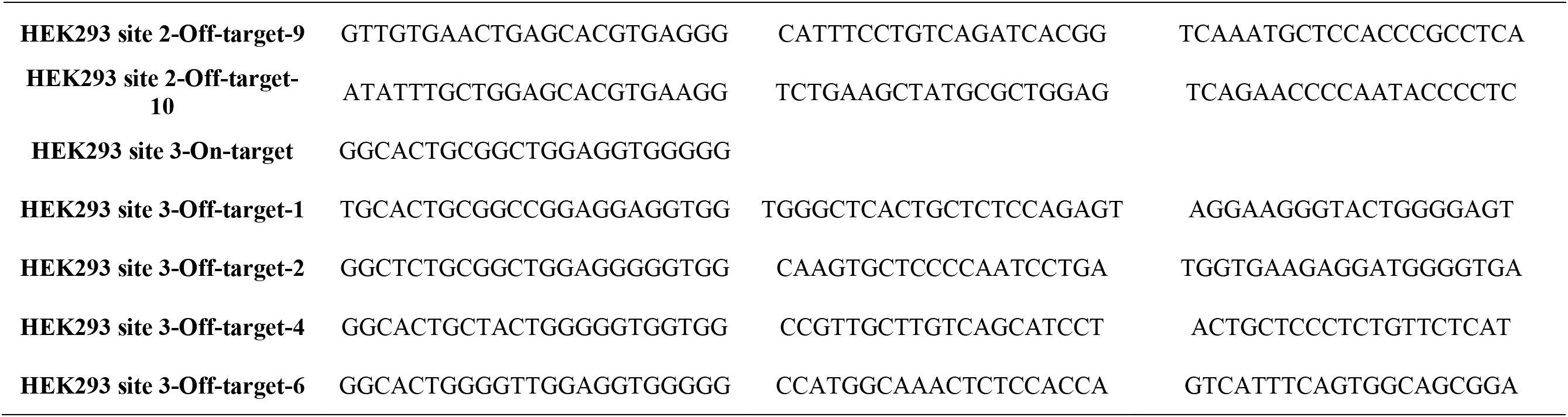
Primers used for deep sequencing of off-target effects.

